# Longitudinal Intravital Imaging of Biosensor-labeled *In Situ* Islet Beta Cells

**DOI:** 10.1101/518753

**Authors:** Christopher A. Reissaus, Ashley N. Twigg, Kara S. Orr, Abass M. Conteh, Michelle M. Martinez, Malgorzata M. Kamocka, Richard N. Day, Sarah A. Tersey, Kenneth W. Dunn, Raghavendra G. Mirmira, Amelia K. Linnemann

## Abstract

Impaired function and apoptosis of insulin-secreting islet β-cells is central to disease progression in both type 1 and type 2 diabetes. Oxidative damage resulting from excess reactive oxygen species (ROS) is a central factor in β-cell dysfunction and death, but the dynamic nature of ROS accumulation and its depletion pose a problem for mechanistic studies *in vivo*. Biosensors, including the redox-sensitive GFP (roGFPs), coupled with intravital microscopy provide a sensitive and dynamic solution to this problem. Here, we utilize a virally-delivered roGFP2-containing human glutaredoxin-1 (Grx1-roGFP2) to selectively monitor β-cell ROS dynamics *in vivo* in response to toxic glucose analogs. We paired viral biosensor delivery with implanted abdominal imaging windows over the pancreas, thus allowing longitudinal measurements of β-cell ROS and islet area during and after streptozotocin (STZ) exposure. The studies presented here represent a robust experimental platform that could be readily adapted to various transgenic or physiological mouse models in conjunction with any number of available biosensors, and thus opens a vast realm of potential for discovery in islet biology *in vivo*.

## Introduction

The Islet of Langerhans is a complex micro-organ composed of multiple cell types, including neuro-endocrine, endothelial, neuronal, and immune cells. These cells work in concert to establish a tightly-regulated milieu that plays a central role in glucose homeostasis. This unique environment is lost when islets are isolated for characterization *in vitro*. While some of the *in vivo* characteristics of islets can be recovered using novel culture methods and engineered microenvironments^1^, more targeted approaches utilizing intravital microscopy have yielded crucial data regarding islet biology *in situ*^2,3^. However, intravital studies in islet biology are currently limited because of the complexity of the experimental regime and the lack of easily implemented tools to target specific cells for analysis of physiological function. Here, we present methods to overcome these limitations using selective β-cell expression of a virally packaged biosensor, Grx1-roGFP2, to measure the dynamics of ROS regulation *in vivo*.

Oxidative stress occurs when increased production of cellular oxidants (e.g. ROS) is not balanced by an increase in cellular antioxidants/antioxidant enzymes. A considerable body of evidence supports the conclusion that oxidative stress is a common feature of both type 1 and 2 diabetes (T1D/T2D)^4,5^. When unresolved, oxidative stress can trigger β-cell apoptosis. The ability to study these processes in intact islets within their native environment is currently limited by the lack of sensitive real-time sensors of ROS generation that can be delivered to specific cells *in vivo*. To address this, we developed a platform that simplifies the introduction of a cell-specific biosensor without the need for generation or breeding of biosensor transgenic animals, then analyzed the biosensor’s signaling both *in vitro* and *in vivo*. To enable longitudinal imaging, we coupled this platform with abdominal imaging windows. This allowed us to make the first observations of β-cell specific ROS responses over the time-course of STZ exposure, and to couple our observations of progressive β-cell loss with the development of hyperglycemia.

## Results

Several variants of redox-sensitive GFPs, or roGFPs, have been engineered that are sensitive to different parts of the ROS response cascade or within different cellular compartments. Here, we used roGFP2 containing human glutaredoxin-1 (Grx1-roGFP2), which is sensitive to the glutathione redox cycle in the cytoplasm^6^. Though ROS is primarily generated in the mitochondria, literature supports the idea that the unregulated release of mitochondrial ROS into the cytosol and enhanced cytoplasmic production of ROS are key features in the progression of cellular oxidative stress^7,8^. We first generated an adenovirus containing Grx1-roGFP2 under the control of a hybrid insulin/rabbit beta globin promoter^9^ (Ad-lNS-Grx1-roGFP2) to label and selectively measure ROS dynamics in islet β-cells. The specificity of this promoter was confirmed by imaging of adenovirus-infected isolated islets from mice that were transgenic for a β-cell specific nuclear mCherry tag^10^, as well as by immunofluorescence against insulin and GFP in infected isolated islet cells and paraffin-embedded pancreas (Fig S1A-C). Using the β-cell-specific Ad-lNS-Grx1-roGFP2, we first validated biosensor activity in isolated wild-type (WT) C57Bl/6J mouse islets using confocal microscopy. We hypothesized that β-cells might have altered ROS levels upon acute exposure to different glucose concentrations. The short-term elevation (16.7mM) or reduction (2mM) of glucose from basal stimulatory levels (defined here as 8mM) over 30 minutes did not alter the 405nm/488nm ratiometric excitation readout of the biosensor acutely (Fig S1D). We extended incubations to 4 and 16 hours at 2mM, 8mM, 16.7mM, and 25mM glucose levels to exacerbate changes in ROS levels, which resulted in an inverse relationship between glucose concentrations and ROS, as measured by the ratiometric intensities from Grx1-roGFP2 (Figure 1A-C). Next, we treated the biosensor-labeled isolated islets *in vitro* with 4mM alloxan monohydrate, a toxic glucose analog that produces ROS when metabolized, to test whether acute changes in β-cell ROS were measurable with Grx1-roGFP2 (Figure 1D). The magnitude of change in ROS (1.4 fold at 8mM vs. 2.2 fold at 16.7mM) appeared to be related, in part, to the concentration of glucose and the roGFP2 ratio at baseline (Figure 1E). Together, these data demonstrate the ability of roGFP2 to monitor acute and chronic changes in ROS within pancreatic β-cells.

**Figure 1:**
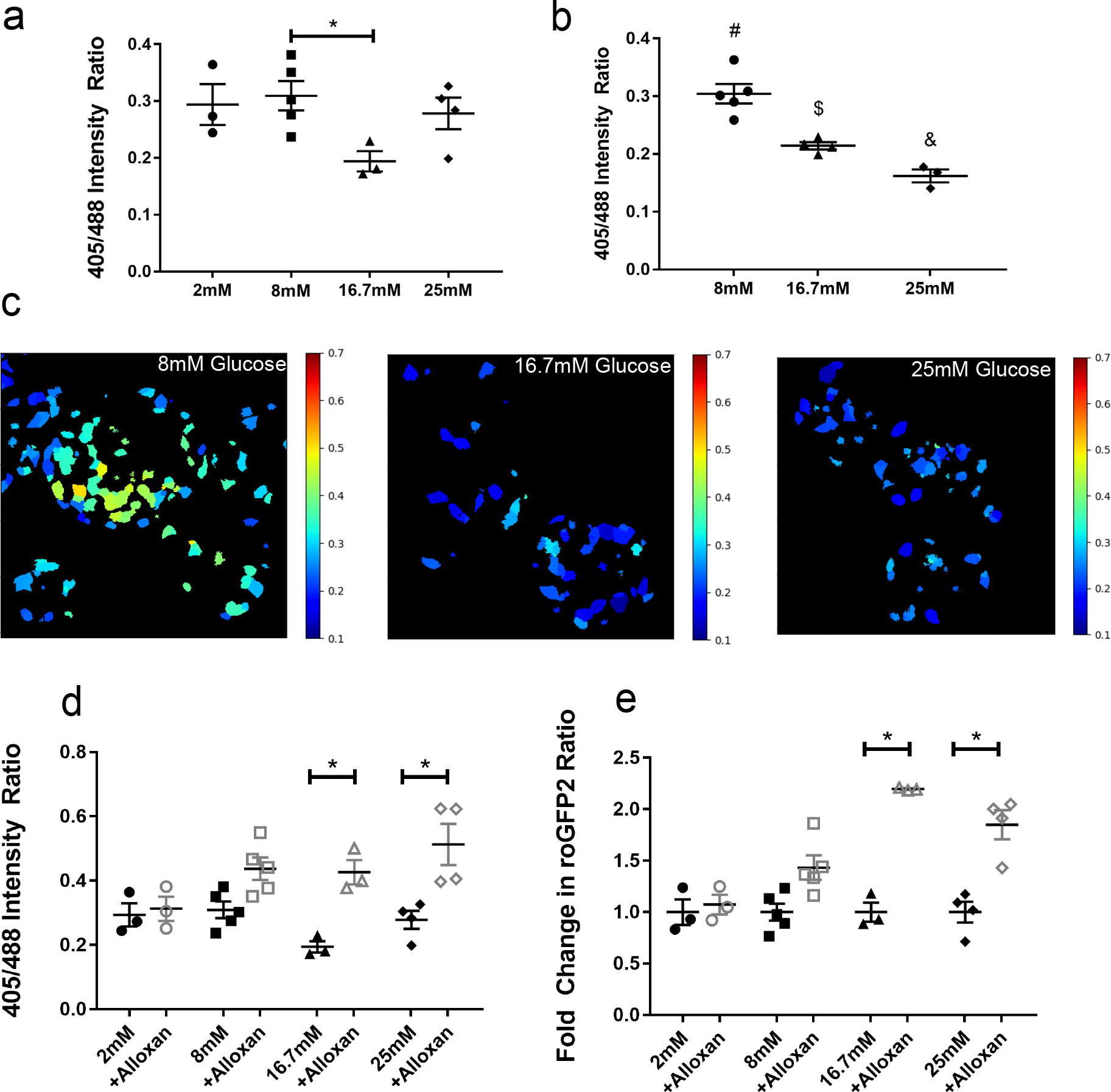
In Vitro Characterization of Adeno-INS-Grx1-roGFP2. (A) Adeno-INS-Grx1-roGFP2 infected primary islets from C57Bl/6J mice were incubated for 4 hours with islet media containing 2, 8, 16.7, or 25 mM glucose. Emissions (490-600nm) were collected from sequential 405 and 488 nm excitation. (B) Similar to A, except islets were incubated for 16 hours with islet media containing 2, 8, 16.7, or 25 mM glucose then imaged. (C) Representative ratiometric images generated from islets after 16 hours glucose incubations. (D) Adeno-INS-Grx1-roGFP2 infected mouse islets incubated for 4 hours with islet media containing 2, 8, 16.7, or 25 mM glucose (filled black) were then imaged after 5 minutes of stimulation with 4mM alloxan monohydrate (hollow grey). (E) Data from (D) were replotted as a fold change relative to the baseline ratio for each mouse. Data are means ±SEM. (N ≥ 3 mice; >4 islets per mouse; p<0.05).

Primary islets maintained *in vitro* are removed from the normal physiological cues once isolated from the organism. Thus, it is critical to develop a platform to monitor cellular changes in the native environment of the intact tissue to fully understand cellular responses and disease progression. To explore the dynamics of ROS in β-cells *in vivo*, we pursued two methodologies to label and measure β-cell-specific roGFP2 signaling using intravital microscopy (Fig 2A). We utilized a syngeneic transplantation model, in which wild-type C57Bl/6J mouse islets infected *in vitro* with Ad-INS-Grx1-roGFP2 were transplanted under the kidney capsule of recipient mice. Once the labeled islets were identified under the kidney capsule, the mice were intravenously (IV) injected with saline, followed by 80mg/kg alloxan monohydrate in saline (Figure 2B). Infusion of saline alone did not alter roGFP2 signal, whereas infusion of alloxan resulted in a 2-fold increase in the roGFP2 ratio within 5 minutes. The signal then decreased over the next 15 minutes, presumably due to activation of the endogenous antioxidant response and mitigation of intracellular ROS within the transplanted islets.

**Figure 2:**
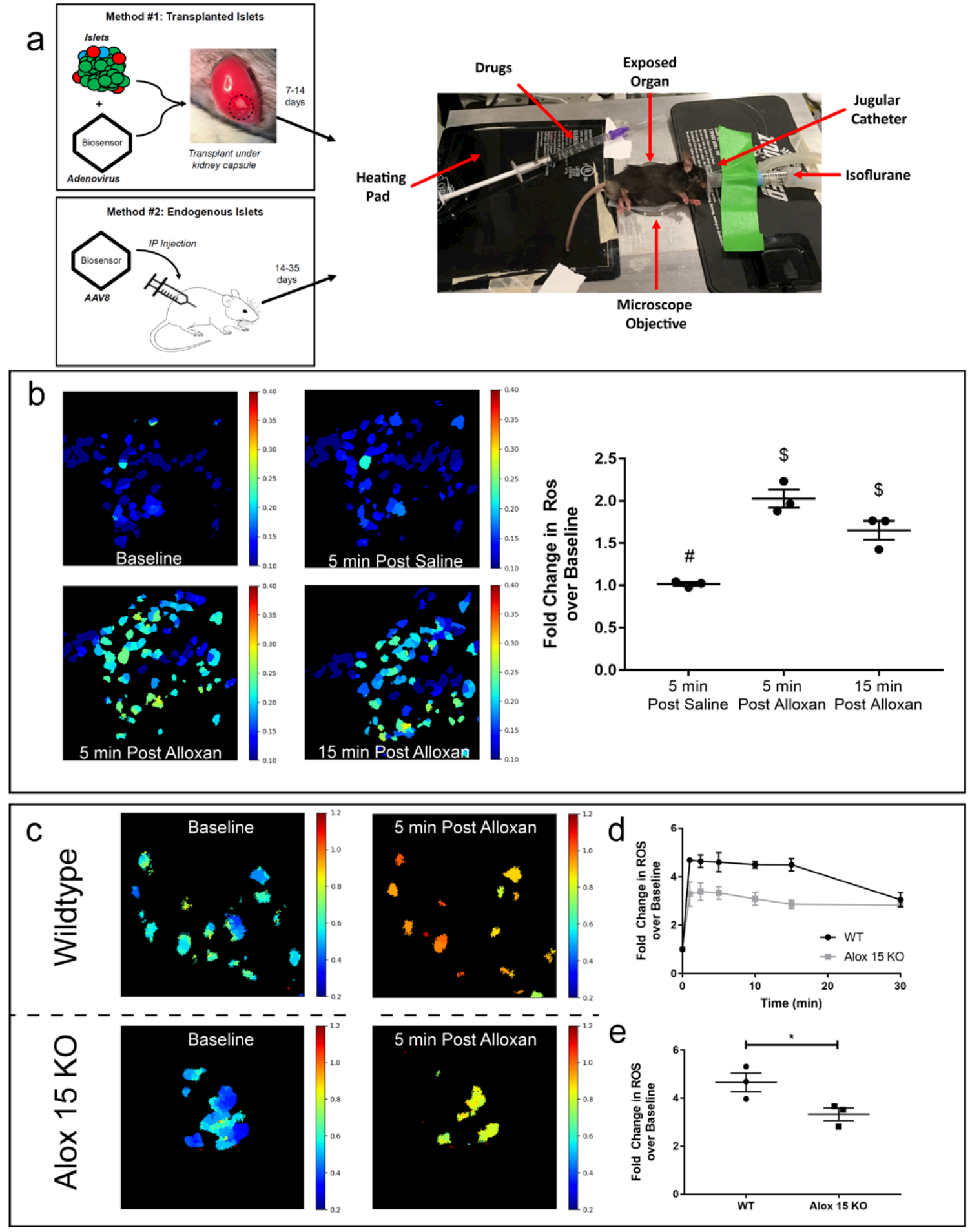
Intravital Microscopy Applications of Adeno- and AAV8-INS-Grx1-roGFP2. (A) Schematic depiction of the two methodologies for imaging of biosensor-labeled β-cells using intravital microscopy. Method #1 utilizes adenoviral labeling of isolated primary mouse islets, followed by transplantation under the mouse kidney capsule. Method #2 uses intraperitoneal (IP) injection of adeno-associated virus serotype 8 (AAV8) packaged biosensor to label endogenous β-cells. Mice are anesthetized using continuous isoflurane inhalation and placed on a heated stage where the imaging objective reaches the exposed kidney or pancreas from below. A jugular catheter is used to administer drugs intravenously. (B) Representative ratiometric images of Grx1-roGPF2 changes in β-cells within the islet transplant under the kidney capsule at baseline, 5 minutes after IV saline injection, and 5 and 15 minutes after IV injection of 80 mg/kg alloxan monohydrate. (C) Fold change over baseline in β-cell-specific Grx1-roGFP2 from 3 mice after 5 minutes after IV saline injection, and 5 and 15 minutes after IV injection of 80 mg/kg alloxan monohydrate. Data are means ±SEM. (N=3, p<0.05). (D) Representative ratiometric images of Grx1-roGPF2 changes in β-cells of WT and *Alox15*^−/−^ mice within the endogenous pancreas at 5 minutes after IV injection of 85 mg/kg alloxan monohydrate. (E) Fold change over baseline in Grx1-roGFP2 signal from *in situ* β-cells in the pancreata of WT (black) and *Alox15*^−/−^ (grey) mice over 30 minutes post IV injection of alloxan. (F) Fold change over baseline in Grx1-roGFP2 signal at 5 minutes post IV injection of alloxan.

While these studies demonstrated the utility of the Grx1-roGFP2 biosensor in transplanted islets, we also wanted to develop a system to measure β-cell function *in situ* in the pancreas. To achieve this objective, we developed an adeno-associated virus serotype 8 (AAV8) containing INS-Grx1-roGFP2 (AAV8-INS-Grx1-roGFP2). AAV8 has pancreas tropism^11^, and when combined with the insulin promoter, the biosensor can be selectively expressed in endogenous β-cells (Fig S1A/B). In practice, the AAV8 could be injected into any animal model to be used for intravital imaging, without the need for lengthy breeding of transgenic mice expressing the biosensor in islet tissue. Therefore, to test the portability of our AAV8-packaged biosensor, we used a mouse model that would confer altered antioxidant response compared to WT mice. *Alox15* knockout mice (*Alox15*^−/−^) lack the 12/15-lipoxgenase enzyme, which we previously reported to confer protection against oxidative stress^12^. We hypothesized that this animal model would therefore exhibit altered kinetics of ROS dynamics in the β-cell that would correlate with changes in oxidative damage compared to WT controls. *Alox15*^−/−^ and WT littermates were each administered a single intraperitoneal (IP) injection of AAV8-INS-Grx1-roGFP2 three weeks prior to intravital imaging to allow for biosensor expression. For imaging of ROS dynamics *in vivo*, labeled superficial islets were identified in the intact pancreas. The mice then received IV infusions of 80mg/kg alloxan. Consistent with our expectations, we observed a rapid 4.6-fold increase in biosensor signal ratio in WT mice, compared to a 3.3-fold increase in *Alox15*^−/−^ mice (Fig 2C). Together, these data demonstrate that the intravital imaging of both β-cells transplanted under the kidney capsule and β-cells within the intact pancreas can be used to measure β-cell ROS dynamics *in vivo*.

While acute ROS dynamics can be predictive for β-cell damage, chronic changes leading to oxidative stress are more prominent during the pathogenesis of diabetes. To capture chronic, longitudinal changes, we implanted a modified version of a previously described abdominal imaging window (AIW) into WT mice^13^. AIWs provide a stable method to image the endogenous tissue on the time scale of days to weeks^13^, allowing stable imaging of the same region of pancreas over time. Thus, we used AIWs in AAV8-INS-Grx1-roGFP2-injected WT mice to visualize and measure longitudinal changes in β-cell ROS, as well as β-cell mass after exposure to the selective β-cell toxin streptozotocin (STZ) (Fig 3A). Animals were administered 5 daily, single injections of 55 mg/kg STZ (the multiple low-dose STZ, or MLD-STZ challenge). MLD-STZ is a well-characterized pharmacological model to induce diabetes, where hyperglycemia develops within days after the last STZ injection, and is associated with reduced β-cell mass due to apoptosis^14,15^. Figure 3B shows an example of a pancreas window preparation, demonstrating that stable positioning is obtained within ~8 days of window placement, and is maintained for an additional 24 days. The addition of STZ did not cause a noticeable change to the overall appearance of the tissue for the duration of longitudinal imaging. Baseline microscopy images were collected 11 days after window implantation, prior to beginning the MLD-STZ challenge (Fig 3C). STZ injections were administered in the afternoon, while the imaging was completed in the morning to avoid measuring acute effects of the drug, but instead to measure the chronic effects on ROS that persist over time. Images were collected sequentially from the same islet before, during, and up to 17 days after MLD-STZ (Fig 3C).

**Figure 3:**
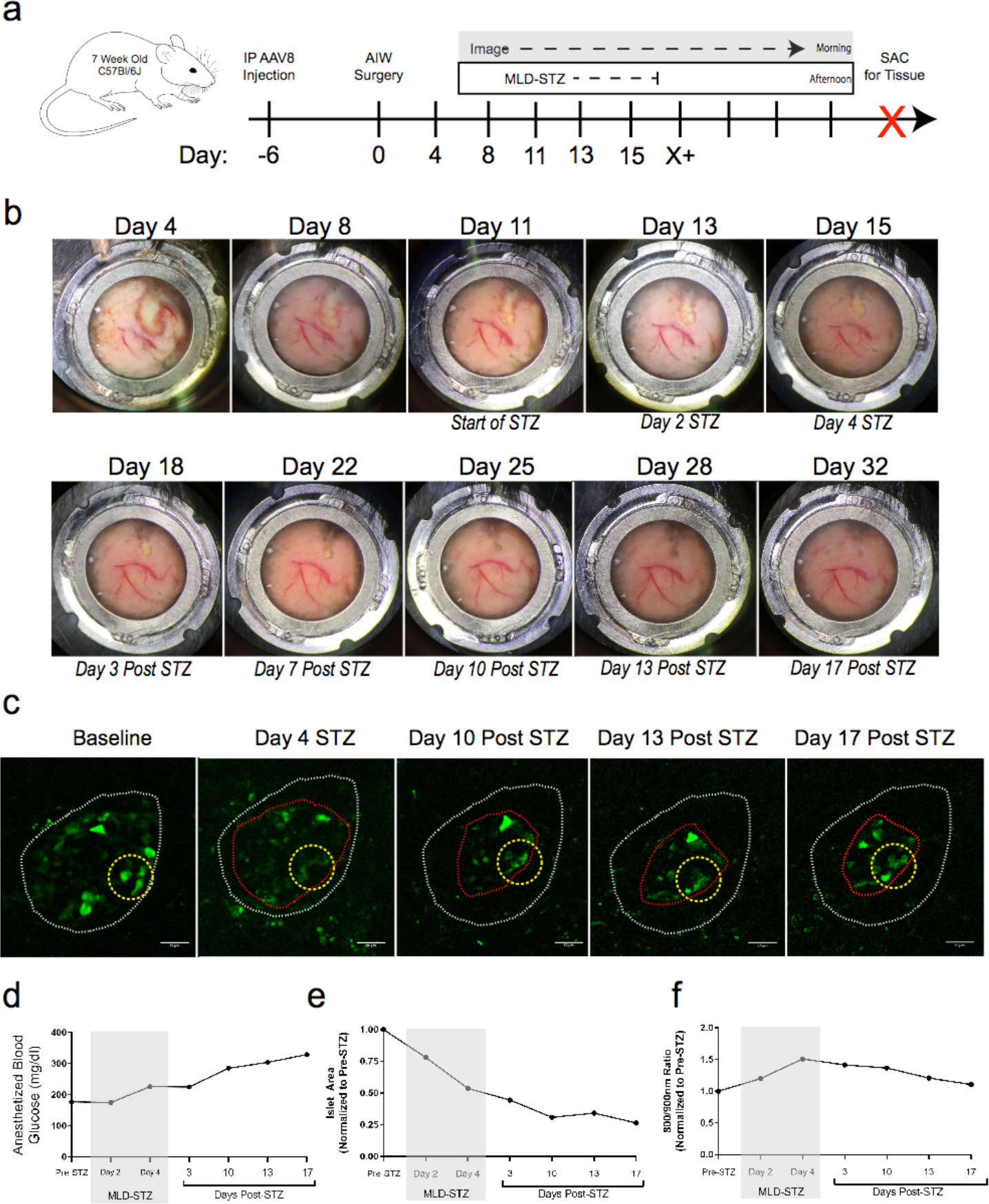
Longitudinal Imaging of AAV8-INS-Grx1-roGFP2 using Abdominal Imaging Windows after Multi-low Dose STZ. (A) Schematic of AAV8-INS-Grx1-roGFP2, abdominal imaging window (AIW), and multi-low-dose streptozotocin (MLD-STZ; 55mg/kg/day) experiment. Intraperitoneal injection of AAV8-packaged biosensor occurred 6 days prior to AIW surgery (day 0). Baseline images were collected in the morning of day 11 and MLD-STZ started in the afternoon of day 11. MLD-STZ continued for 5 days while imaging continued until the window integrity was compromised at Day 32. The pancreas was recovered and fixed for endpoint analyses. (B) Representative widefield images of the pancreas as seen through the AIW over time. Above each image is the day post AIW implantation. Below each image is the corresponding day in relation to MLD-STZ challenge. (C) Representative images of a single z-plane within an imaged islet at baseline, day 4 of MLD-STZ, day 10 post-STZ, day 13 post-STZ, and day 17 post-STZ. The white dotted line indicates the largest x/y islet area at baseline. The yellow dotted line highlights a similar region of moderately stable cells that aided in aligning z-planes over time. The red dotted line shows the final, largest x/y islet area 17 days after MLD-STZ challenge. (D) Representative blood glucose collected 30 minutes after start of anesthesia for each imaging session. (E) Islet area calculated for the largest x/y plane (black dotted line) before, during, and after MLD-STZ challenge. (F) Ratiometric data (black dotted line) from AAV8-INS-Grx1-roGFP2-labeled β-cells using 800/900 nm 2-photon excitation before, during, and after MLD-STZ challenge.

Representative z-planes demonstrate that while the same islet was imaged over time, the islet architecture changed dramatically in response to STZ. We used second harmonic generation from collagen to verify that we were returning to the same islet during each imaging session (Fig S2). Consistent with previous reports^14,15^, MLD-STZ resulted in elevated blood glucose levels by 3 days post-treatment and overt hyperglycemia was observed by 17 days post treatment (Fig 3D). We measured the largest x/y area and observed a rapid 50% reduction in the islet area by the end of STZ treatment, which then stabilized at ~35% of the initial area by 10-days post-treatment (Fig 3E). At the end of intravital studies, animals were euthanized and pancreata were fixed for additional end-point analysis using immunohistochemistry and immunofluorescence to confirm loss of islet area (Fig S3).

The reduced islet area and increased blood glucose levels after MLD-STZ are indicative of β-cell ablation. In order to collect data from islets through the AIW deep within the pancreas, we utilized multi-photon microscopy and sequential 800nm and 900nm excitation of Grx1-roGFP2. While the dynamic range of of roGFPs is less using two-photon excitation^16^, Grx1-roGFP2 ratios measured from the imaged β-cells show that there is an elevated biosensor signal during STZ administration (Fig 3F). This effect subsides ~13 days after MLD-STZ has ended and occurs in conjunction with the stabilization of islet area. The somewhat muted response of the roGFP2 sensor is likely to reflect the fact that measurements are obtained from the subpopulation of cells that have been able to resist STZ ablation. We have replicated these results in additional mice to demonstrate the uniformity of the STZ ablation compared to unchanging saline-treated controls (Fig 4).

**Figure 4:**
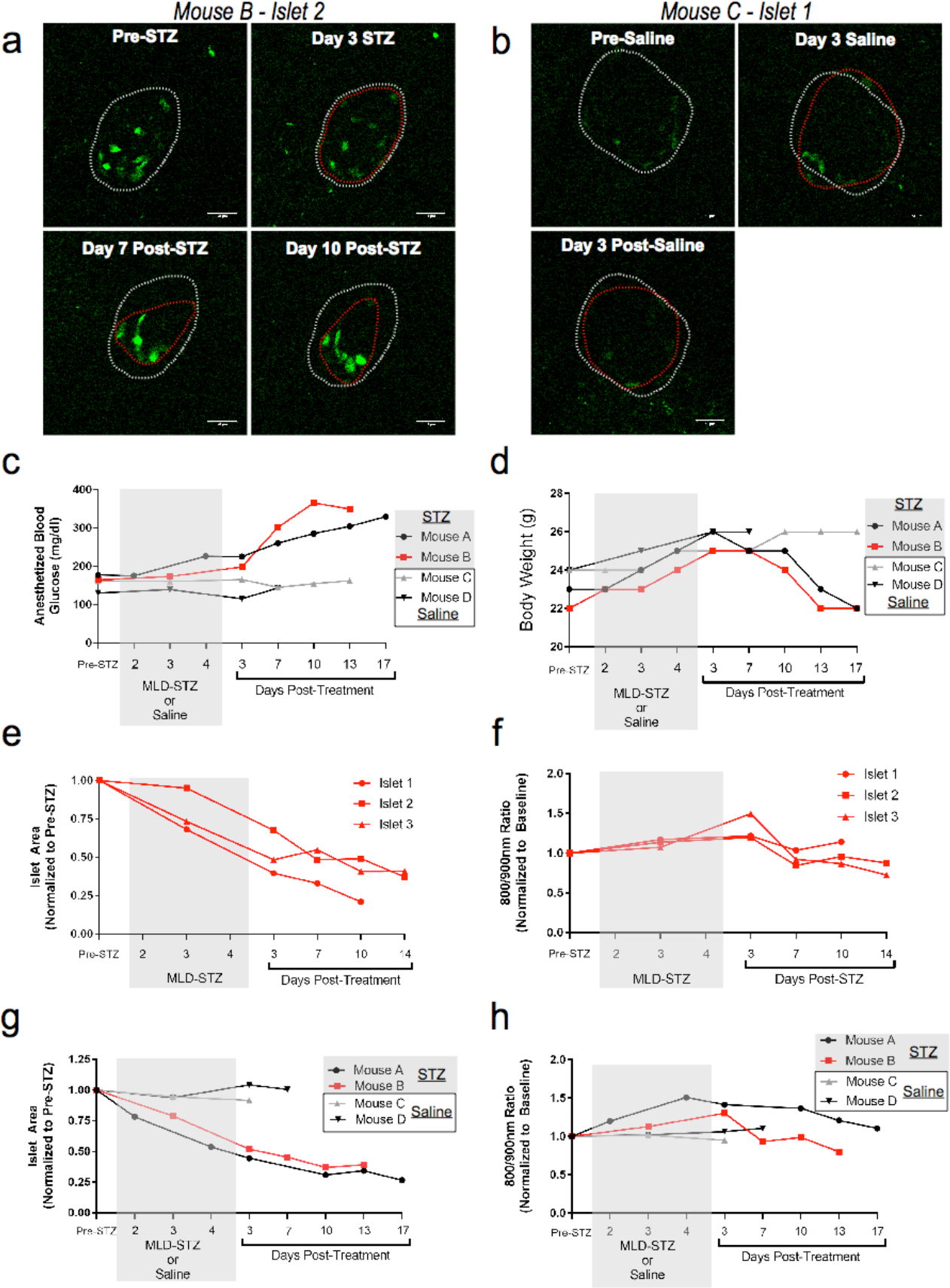
Longitudinal Imaging of AAV8-INS-Grx1-roGFP2 using Abdominal Imaging Windows after Multi-low Dose STZ or Saline. (A) Representative slices from z-stacks of the same islet pre-STZ, 3 days during STZ treatment, 7 and 10 days post-STZ treatment for Mouse B. The white dotted light highlights the starting islet area, while the red dotted line shows the islet area of that day’s imaging. (B) Representative slices from z-stacks of the same islet pre-saline, 3 days during Saline treatment, and 3 days post-saline treatment for Mouse C. The white dotted light highlights the starting islet area, while the red dotted line shows the islet area of that day’s imaging. (C) Blood glucose was collected 30 minutes into anesthesia for each imaging session. Mouse A (black circles) and B (red squares) received STZ, while Mouse C (gray triangles) and D (black triangles) received saline injections. (D) Body weight (in grams) of mice that had undergone AIW study. (E) Islet area calculated for the largest x/y plane for 3 individual islets for Mouse B before, during, and after MLD-STZ challenge. (F) Ratiometric data the 3 individual islets from Mouse B using AAV8-INS-Grx1-roGFP2-labeled β-cells after 800/900 nm 2-photon excitation before, during, and after MLD-STZ challenge. (G) Aggregated islet area data from Mouse A (black circles), B (red squares), C (gray triangles), and D (black triangles) before, during, and after STZ or saline injections. (H) Ratiometric data from Mouse A (black circles), B (red squares), C (gray triangles), and D (black triangles) before, during, and after STZ or saline injections in AAV8-INS-Grx1-roGFP2-labeled β-cells after 800/900 nm 2-photon excitation before, during, and after MLD-STZ challenge.

## Discussion

We have demonstrated the application of a ROS biosensor within the context of β-cell biology. The use of Grx1-roGFP2 in β-cells both *in vitro* and *in vivo* enables acute and chronic measurements of ROS regulation. These data offer a stepping stone to more complex analysis focused on determining why specific cell populations and subpopulations are more susceptible to ROS. With the ability to measure acute ROS dynamics, we may also begin to tease apart inter- and intra-islet differences in oxidative stress susceptibility between mouse and human β-cells ^17^. While the pathophysiology of β-cell oxidative stress continues to be elucidated, other known signaling pathways, including calcium signaling, are also critical to β-cell function and survival in diabetes^18^. With this in mind, the adeno- and AAV-applications have been designed to permit the easy substitution of biosensors for other pathways of interest (e.g. GCaMP or Twitch for calcium). Furthermore, these virally-packaged biosensors can be utilized in virtually any mouse model without lengthy breeding, or may also be used in human islets transplanted into mice.

While there are moderate differences in alloxan stimulation between transplanted and endogenous β-cells likely caused by the stress of transplantation and/or efficiency of graft vascularization, our intravital microscopy data from biosensor-labeled β-cells demonstrate the importance of acute dynamics in cellular responses to stimuli. However, chronic changes in oxidative stress may be more indicative of normal diabetes progression. With the use of AIW’s, we demonstrate that longitudinal imaging of biosensor-labeled β-cells can yield a robust data set, including both physiological and cellular measurements, which can be chronologically interpreted to generate a sequence of events during STZ-induced diabetes. Here we are able to observe that blood glucose levels inversely correlated with islet area and that the glucose values are delayed in chronology, suggesting that remaining β-cells are partially able to compensate for the loss in mass up to a “tipping point”^19^. This experimental platform has yielded an unprecedented data set of longitudinal cellular measurements collected from biosensor-labeled β-cells *in situ* during β-cell death. This approach could also easily be adapted to more targeted models of diabetes such as the *db/db* or non-obese diabetic (NOD) mice. In conclusion, our highly-portable experimental platform, comprised of virally-packaged biosensor and AIW's, thus facilitate the unparalleled interrogation of *in vivo* β-cell biology. The longitudinal approach truly opens a new realm of *in vivo* experimentation in islet structure and function within the context of steady-state drug treatments, β-cell replication or death, and diabetes progression.

## Methods

### Islet Isolation

Male and female C57BL/6J mice were purchased from Jackson Labs at 8 weeks of age and utilized in both *in vitro* assays and islet transplant studies at 8-14 weeks of age. 20 week-old female mice expressing an insulin-promoter driven ROSA26 nuclear H2B-mCherry^10^ were utilized for mCherry colocalization studies. Mice were euthanized, pancreas harvested, and islets were liberated using a 0.3% collagenase digestion in 37°C shaking water bath in Hanks buffered sodium salt. Islets were maintained in islet media containing phenol free RPMI 1640, with 10% FBS, 100U/ml Penicillin, 100ug/ml Streptomycin, and 8mM glucose. Islets were allowed to recover over-night prior to any subsequent experiments.

### Virus Generation

Adeno- and adeno-associated viruses (AAV) were generated using VectorBuilder (Cyagen Biosciences). Briefly, the custom β-cell specific promoter contained 414 base pairs of the rat insulin-1 promoter, along with 691 basepairs of the rabbit beta-globin intron^9^. This promoter and Grx1-roGFP2^6^ sequences were de novo synthesized and cloned into the necessary adeno- or AAV-expression vectors for viral production.

### *In Vitro* Islet Infection, Treatment and Imaging

To infect β-cells *in vitro*, isolated islets were washed with Dulbecco’s phosphate buffered saline without calcium or magnesium, followed by distention with Accutase (Sigma) at 37°C for 30 seconds. Accutase was rapidly inactivated with room temperature islet media, then washed with fresh islet media. Adenovirus was added directly to islet media to achieve approximately 2×10^7^ viral particles per 100 islets. Viral infection lasted at least 6 hours prior to image or transplant the following day. After viral infection, islets were placed in fresh islet media with varying glucose concentrations (2, 8, 16.7, and 25 mM) for either 4 or 16 (overnight) hours. Islets were imaged *in vitro* imaging using a Zeiss LSM 800 (40×/1.2 NA/Water Objective) equipped with an Ibidi Stage Top Incubation system (5% CO2, 37°C, 85% Humidity). Grx1-roGFP2 was sequentially excited with 405 and 488 nm lasers at a 3:1 power ratio, with emissions collected from 490-600 nm. Alloxan monohydrate (Sigma) was resuspended in islet media at 4mM immediately prior to the start of imaging.

### Islet Transplants

Adeno-infected islets were transplanted into anesthetized recipient mice via sub-renal capsular injection under aseptic conditions. Mice were anesthetized with isoflurane. The left lumbar region received a single skin incision to expose the kidney. A small entry hole (~1mm diameter) was made in the kidney capsule with a jeweler forceps and islets were gently released into the subcapsular space through a small piece of sterile polyethylene tubing attached to a syringe. The renal capsule was allowed to close by secondary intention, the body wall and skin incision were closed using monofilament 4.0 silk sutures, and the mice were allowed to recover. Analgesics were administered for pain relief. The mice were singly housed post-surgery. Islet transplants were allowed to engraft 14 days prior to intravital imaging.

### End-point Kidney Transplant and Pancreas Intravital Imaging

Mice for kidney capsule islet transplant or pancreas imaging were anesthetized with isoflurane on the day of imaging. The mouse was placed in the right lateral decubitus position and a small, 1 cm vertical incision was made along the left flank at the level of the left kidney or over the pancreas. The exposed organ was orientated so that the organ was underneath the animal for imaging with an inverted Leica SP8 confocal microscope. The animal’s temperature was maintained via a circulating water bath blanket, a second warmer for the stage and heating elements for the objective to diminish the heat sink at the level of the objective tissue interface. A rectal thermometer was in place to keep a continuous read out on the mouse’s core body temperature. Intravenous access was achieved using the placement of a jugular catheter or by tail vein injection. Images were captured using a Leica SP8 (25×/0.95NA/Water immersion objective) with 405 and 488 nm lasers at an 8:1 power ratio, with emissions collected from 490-600 nm. Alloxan monohydrate (Sigma) was resuspended in sterile saline at 80 mg/kg immediately prior to the IV injection and imaging.

### *In Vivo* Islet Infection

To infect β-cells *in vivo*, AAV8-INS-Grx1-roGFP2 was injected intraperitoneally 14-21 days prior to intravital imaging or 8 days prior to AIW surgery. Mice were given a dose of 2-2.5×10^11^ viral particles.

### Abdominal Imaging Window (AIW) Procedure

A general abdominal imaging window (AIW) procedure has been published previously^13^. We adapted the protocol to be successful over the pancreas. The animals were anesthetized via isoflurane and All procedures were performed under aseptic conditions. Warm sterile saline (1 ml) was injected subcutaneously to aid in fluid replacement post procedure. A vertical left flank incision was performed approximately 1.5cm in length. The pancreas was identified and externalized utilizing cotton tipped applicators. A purse-string suture (non-absorbable monofilament) was placed around the incision through both the fascial and the skin layers. Each suture was approximately 0.5cm apart from the next and at least 0.1cm from the edge of the incision. Cyanoacrylate adhesive was applied to the interior of the underside of the outer ring and then it was placed on the pancreas. Mild pressure was applied for approximately 5 minutes to ensure that the ring adhered to the pancreas. The inner ring with attached coverslip was then placed into the outer ring and contact with the organ of interest was visually confirmed. The skin and fascial edges were then placed into the groove of the AIW and the purse string suture was secured with at least 4 square knots. This effectively closed off the abdominal cavity and left the organ of interest in contact with the coverslip. Once the AIW was in place the mouse was allowed to emerge from anesthesia and placed in a recovery cage on a warming blanket with soft bedding.

### Intravital for AIW Imaging

Mice with AlW’s were anesthetized with isoflurane for no longer than 30 minutes for imaging of labeled islets. Mice received 200uL of sterile saline to prevent dehydration. lmages were captured using a Leica SP8 (25x/0.95NA/Water Objective) equipped with a multiphoton MaiTai Ti-Sapphire 2-photon laser. Z-stacks were sequentially collected using 800 and 900 nm excitation (1:1 power ratio) and a green emission filter (520-580nm). At the end of imaging, duplicated blood glucose readings from a tail nick were measured using an AlphaTrack2 glucometer and test strips. Mice were allowed to recover prior to returning to animal housing.

### Pancreas Recovery, Sectioning, and Staining after AIW

At the end of intravital study, the pancreas was recovered via pancreatectomy and fixed in 4% PFA at room temperature for 4 hours. The pancreas was then dehydrated in 70% ethanol overnight. The pancreas was embedded in paraffin and sectioned for subsequent staining analyses. Pancreata were stained for immunofluorescence using anti-insulin antibody (A0564, Dako; 1;500) and Alexa Fluor 488 anti-Guinea pig antibody (lnvitrogen). lmmunofluorescence images were collected using a Zeiss LSM 700 (40x/1.3NA oil immersion objective). lmmunohistochemistry was performed using anti-insulin antibody (3014, Cell Signaling, 1:400) and anti-Rabbit lmmPRESS HRP reagent kit (MP-7401, Vector labs) then developed using DAB peroxidase Substrate kit (SK-4100, Vector labs). lmmunohistochemistry images were acquired using a Zeiss AxioScan with an Orca ER CCD camera.

### Data Analysis and Image Presentation

Background subtracted, raw images were analyzed using lmageJ (NlH) or Cell Profiler^20^. lntensity of the 405nm-based emission was divided by 488nm-based emission to create a raw Grx1-roGFP2 ratio. Ratiometric images were created using Cell Profiler and rescaled to an arbitrary linear look-up table that fit all data points within that experimental data set. The Cell Profiler Pipeline script is available in the supplemental data. To determine islet area, the x/y plane with the largest GFP+ cell area was outlined and measured using lmageJ. lmages presented in figures were linearly scaled for display purposes only. For statistical testing, Student's t-tests and 1-way ANOVAs were used.

## Acknowledgements

This research was performed using resources and/or funding provided by National Institutes of Health grants K01 DK102492 (to AKL), R03 DK115990 (to AKL), R01 DK60581 and R01 DK105588 (both to RGM), T32 KD0604466 (to CAR), and Human Islet Research Network UC4 DK104162 (to AKL; RRID:SCR_014393), along with startup funds from Indiana University School of Medicine and the Herman B Wells Center for Pediatric Research (to AKL). This study utilized Diabetes Center core resources supported by National Institutes of Health grant P30 DK097512 (to Indiana University). We would like to thank the Indiana O'Brien Center for Advanced Microscopy Analysis for the generous sharing of AIW-related resources. We also thank Mark Soompa for the fabrication of our stage-window adapter and window accessories. We would also like to acknowledge the highly skilled individuals providing services through the Indiana Center for Biological Microscopy and IU School of Medicine Islet Core.

## Contributions

CAR, AT, and AC contributed to data collection and data analysis. CAR, KO, MM, GM, and ST contributed to animal protocols, surgeries, and handling. CAR, RD, ST, KD, RM, and AKL contributed to experimental design, technical oversight, and interpretation of data. CAR and AKL contributed to drafting and writing of the manuscript. All authors contributed to the review and revision of the manuscript.

## Data Availability

The authors declare that all data supporting the findings of this study are available within the paper and its supplementary information files.

---

**Figure S1:**
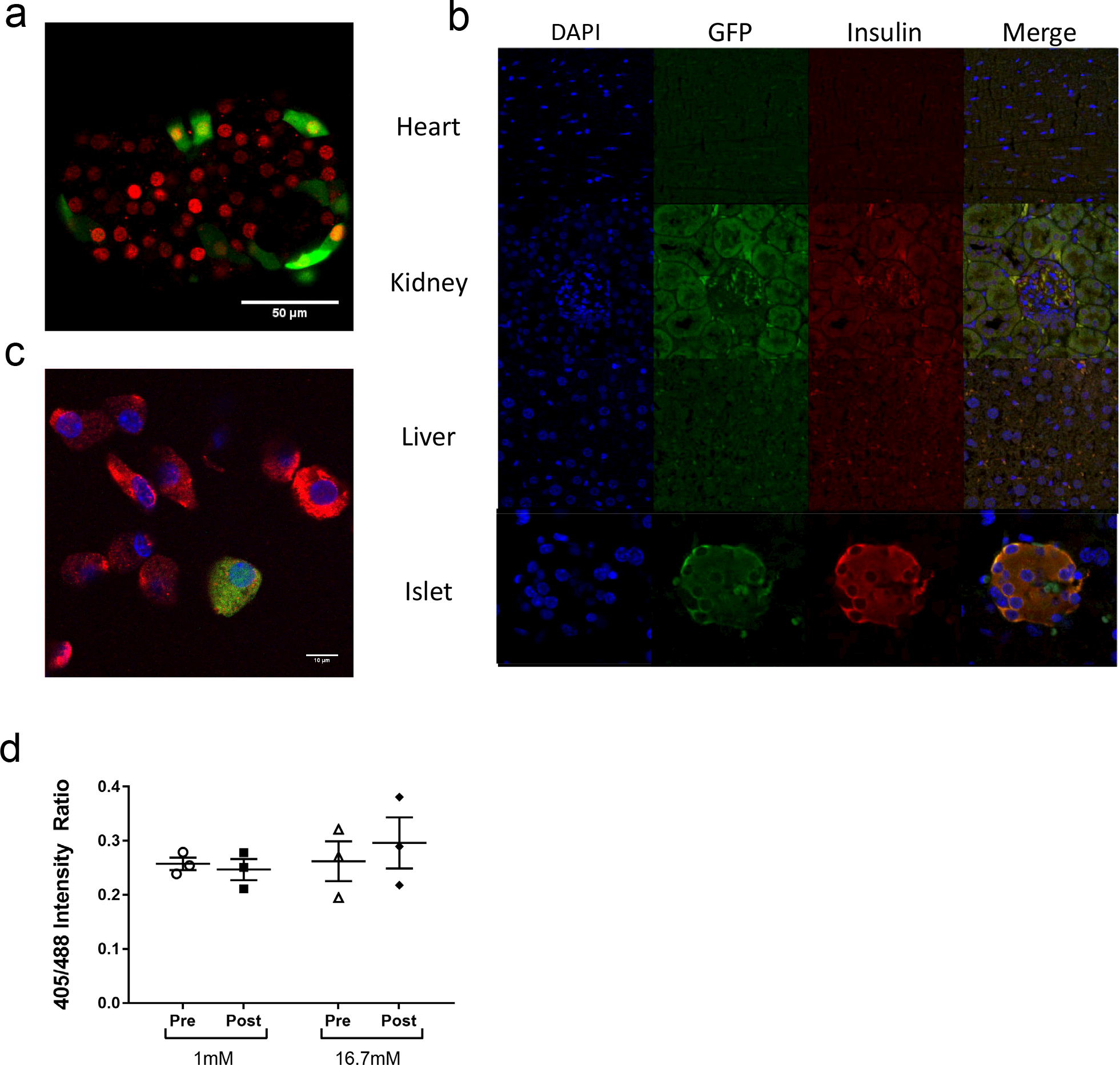
Colocalization of Adeno- and AAv8- IN5-Grx1-roGFP2 *in vitro* and *in vivo*. (A) Isolated islets from mice expressing an insulin-promoter driven R0SA26 nuclear H2B-mCherry were infected with Adeno-INS-Grx1-roGFP2 and imaged using a Zeiss LSM 800 confocal (Green – roGFP2, Red – mCherry). While not all β-cells are infected and express biosensor, all cells that are expressing roGFP2 are mCherry positive. (B) WT mice were IP injected with AAV8-INS-Grx1-roGFP2 3 weeks prior to pancreatectomy, paraffin-embedding, sectioning, and staining. An anti-GFP (to stain roGFP2) and anti-insulin antibodies were used to determine the localization of roGFP2 expression in heart, kidney, liver, and pancreas. (C) Islets were isolated from WT mice IP injected with AAV8-INS-Grx1-roGFP2 3 weeks prior. The islets were dispersed and stained as in (B) and imaged. While AAV8-INS-Grx1-roGFP2 does not label all insulin-positive cells, all GFP-positive cells are also insulin positive. (D) Islets isolated from WT mice were infected with Adeno-INS-Grx1-roGFP2 and treated with 1 and 16.7 mM glucose for 30 minutes. RoGFP2 ratios were collected before and after treatment, however, no changes in biosensor signal were observed. Data are means ±SEM (N=3 mice, >3 islets per mouse).

**Figure S2:**
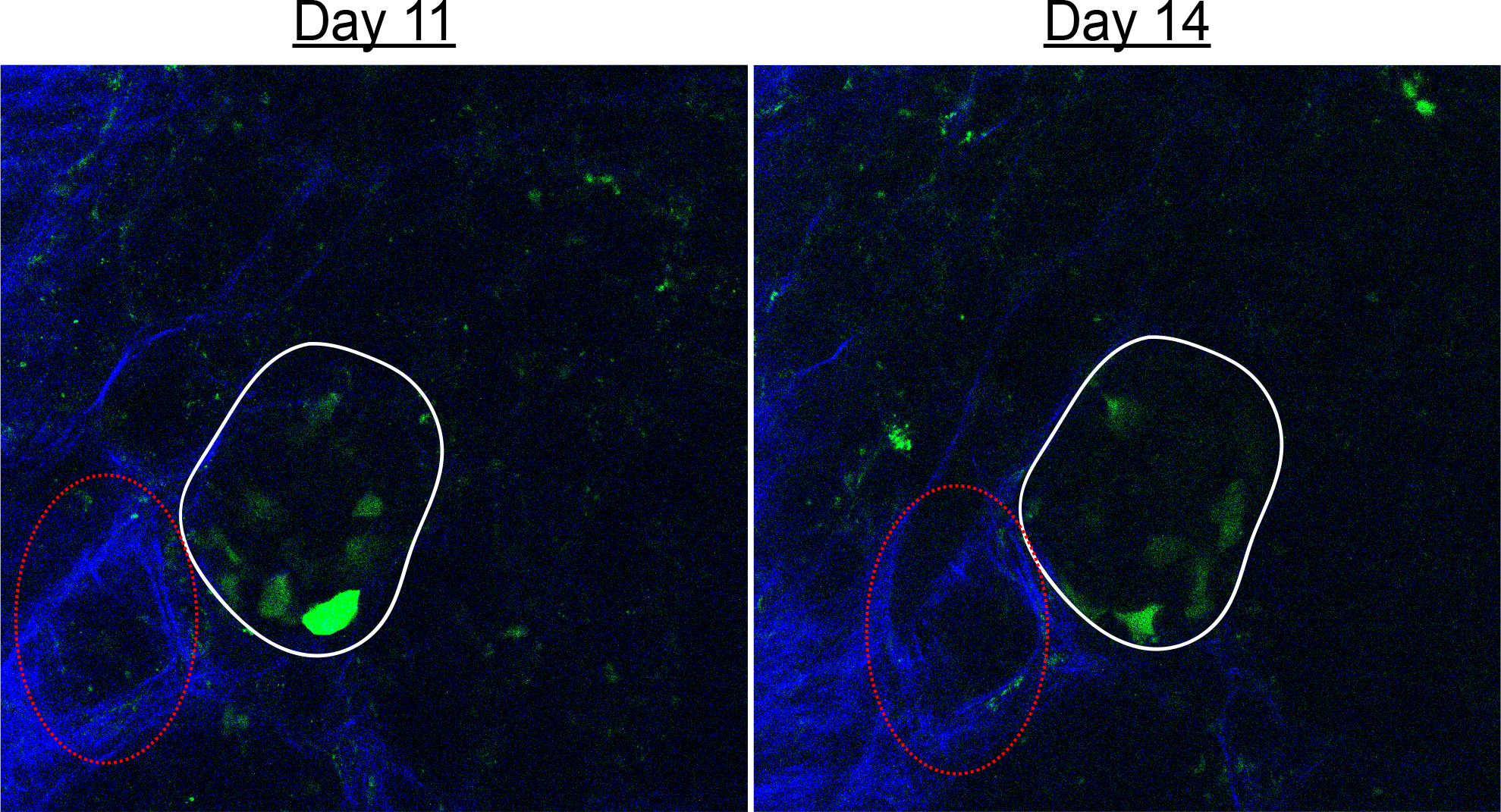
Second Harmonics Act as a Fiduciary Marker. Second harmonic generation from collagen was used as a fiduciary marker to return to islets between imaging sessions. Between days 11 and 14, the islet of interest is highlighted in white, while a unique collagen structure is highlighted in red.

**Figure S3:**
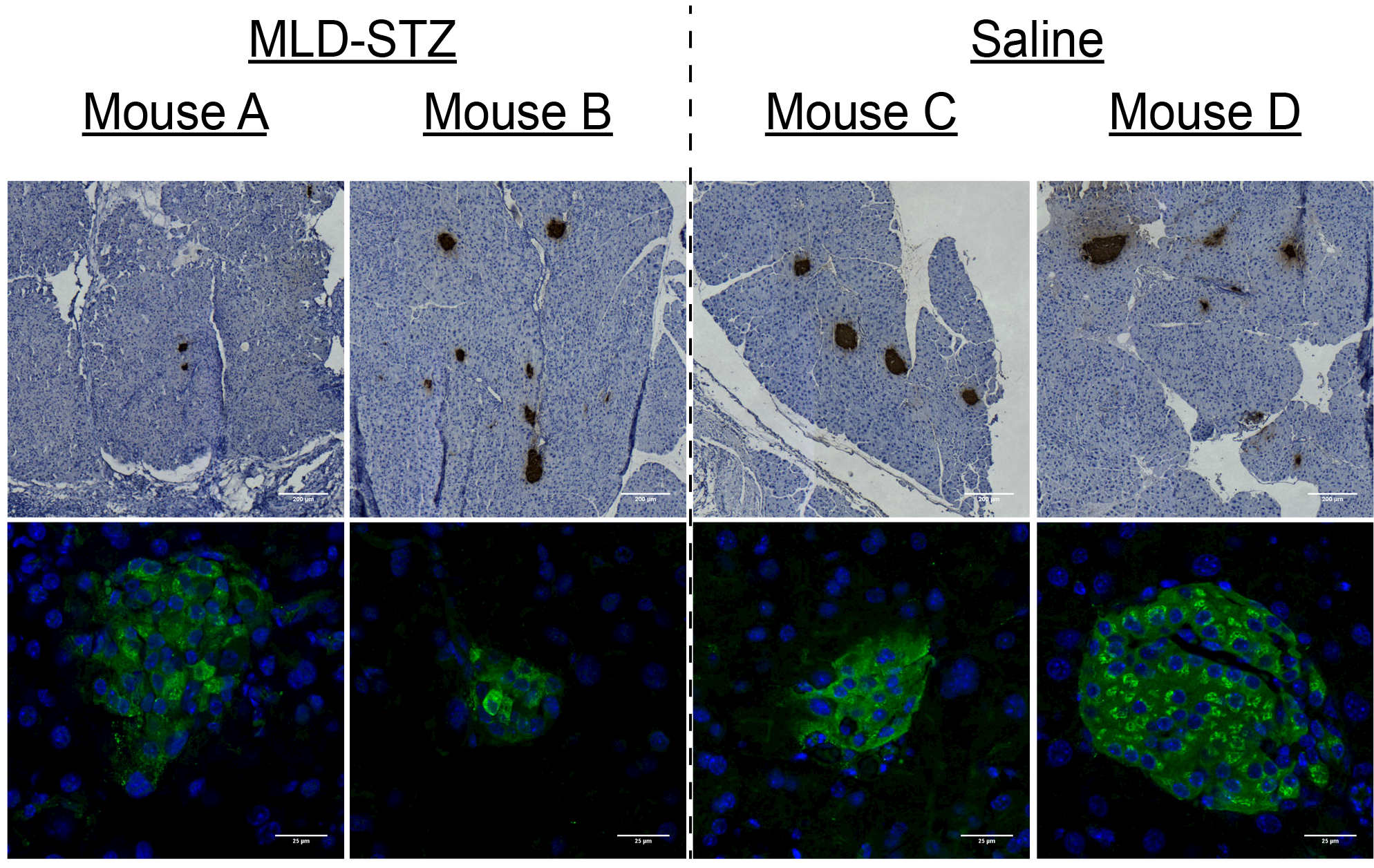
Endpoint Analysis of Pancreata after AIW Study. (A) Representative images of immunohistochemical images of pancreata from the pancreas of Mouse A-D. (B) Representative images of immunofluorescence images (DAPI - Blue, Insulin - Green) of islets from pancreata from Mouse A-D.

